# New insights into *Trypanosoma cruzi* evolution and genotyping based on system-wide protein expression profiles (PhyloQuant)

**DOI:** 10.1101/2020.02.21.959767

**Authors:** Simon Ngao Mule, Andrè Guillherme da Costa Martins, Livia Rosa-Fernandes, Gilberto Santos de Oliveira, Carla Monadeli Rodrigues, Daniel Quina, Graziella E. Rosein, Marta Maria Geraldes Teixeira, Giuseppe Palmisano

## Abstract

The etiological agent of Chagas disease, *Trypanosoma cruzi*, is subdivided into seven genetic subdivisions termed discrete typing units (DTUs), TcI-TcVI and Tcbat. The relevance of *T. cruzi* genetic diversity to the variable clinical course of the disease, virulence, pathogenicity, drug resistance, transmission cycles and ecological distribution justifies the concerted efforts towards understanding the population structure of *T. cruzi* strains. In this study, we introduce a novel approach termed ‘*phyloquant*’ to infer the evolutionary relationships and assignment of *T. cruzi* strains to their DTUs based on differential protein expression profiles evidenced by bottom up large scale mass spectrometry-based quantitative proteomic features. Mass spectrometry features analyzed using parsimony (MS1, iBAQ and LFQ) showed a close correlation between protein expression and *T. cruzi* DTUs and closely related trypanosome species. Although alternative topologies with minor differences between the three MS features analyzed were demonstrated, we show congruence to well accepted evolutionary relationships of *T. cruzi* DTUs; in all analyses TcI and Tcbat were sister groups, and the parental nature of genotype TcII and the hybrid genotypes TcV/TcVI were corroborated. Character mapping of genetic distance matrices based on phylogenetics and phyloquant clustering showed statistically significant correlations. We propose the first quantitative shotgun proteomics approach as a complement strategy to the genetic-based assignment of *T. cruzi* strains to DTUs and evolutionary inferences. Moreover, this approach allows for the identification of differentially regulated and strain/DTU/species-specific proteins, with potential application in the identification of strain/DTU specific biomarkers and candidate therapeutic targets. In addition, the correlation between multi-gene protein expression and divergence of trypanosome species was evaluated, adding another level to understand the genetic subdivisions among *T. cruzi* DTUs.

## Introduction

*Trypanosoma cruzi* is a unicellular protistan parasite agent of Chagas disease (American trypanosomiasis), a zoonotic disease endemic in 21 Latin American countries (1). Here, the World Health Organization (WHO) estimates that approximately 25 million people are at risk of infection, with an estimated 5.7 million infected persons (2). The hematophagous triatomine insects *(Reduviidae: Triatominae)* are the vectors of *T. cruzi*. Other transmission routes include blood transfusion (3–5), oral transmission from food or beverage contamination (6–9), congenital transmission (10, 11) and organ and marrow transplant from infected donors (12, 13). Chagas disease is currently an emergent public health concern with non-vectorial transmission routes and international migrations contributing to sporadic cases of the disease in non-endemic countries including Europe, USA, Australia and Japan (14, 15). Approximately 8 million persons are infected in the world, with 10,000 reported deaths each year (16). During its life cycle, *T. cruzi* alternates between triatomine vectors and mammalian hosts (17). Epimastigotes proliferate in the insect midgut and differentiate into metacyclic trypomastigotes, the non-proliferative infective stage that infect mammals through skin wounds or mucosal membranes, where they invade the cells and differentiate into amastigotes. The amastigotes replicate and differentiate into infective trypomastigotes, which are released into the bloodstream following host cell rapture (18). Each developmental stage is characterized by changes in morphology, molecular and biochemical makeup (17, 19–23). The variable clinical outcomes of Chagas disease have been associated to *T. cruzi* heterogeneity (24–26). Different studies have elucidated the genetic variability in *T. cruzi* populations and shown close associations between the parasites’ genetic and relevant biological, pathological and clinical characteristics (24, 25, 27–29). Currently, six genetic subdivisions, termed discrete typing units (DTUs) are recognized in *T. cruzi* (TcI-TcVI) (30), along with Tcbat which infects predominantly bats (31). The parasite’s DTUs has been implicated as one relevant factor influencing clinical variations of the disease (25, 27, 32), drug resistance/ susceptibility (33, 34), pathogenicity in mice (35), geographic distribution (36), and vector competence (18, 37). Other factors influencing disease outcome include mixed infections, infection routes, host genetics and a range of eco-geographical factors (30, 38). Different molecular techniques have been used to study the genetic diversity of *T. cruzi* such as kinetoplast DNA minicircle (39), RAPDs (40) and more recently by polymorphism of the spliced leader intergenic region (SL-IR) (41–43) and multilocus sequence typing (44–46). In 1999, a consensus nomenclature was proposed, placing *T. cruzi* strains into two major groups; *T. cruzi* I and *T. cruzi* II (47). In 2009, this classification was revised, placing *T. cruzi* genetic subdivisions into 6 discrete typing units (48). In 2012, Tcbat was added as a novel infra-specific genotypes of *T. cruzi* (30). Telleria and colleagues described the hierarchical clustering of *T. cruzi strains* based on gene expression levels from proteomics analysis following 2D-DIGE using Euclidean Distance (49). This clustering showed a high correlation between *T. cruzi* phylogenetics and levels of protein expression. In the current study, we applied a proteome-wide quantitative proteomics to profile and infer evolutionary relationships between *T. cruzi* DTUs (TcI-TcVI, Tcbat) and their most closely related bat trypanosomes; *Trypanosoma cruzi marinkellei, Trypanosoma dionisii, and Trypanosoma* erneyi, all classified in the subgenus *Schizotrypanum* nested into the clade *T. cruzi* (50–52). The generalist and human-infective *T. rangeli* sp. also belonging to the *T. cruzi* clade was included as an outgroup of *Schizotrypanum* (53). The term “phyloquant” is herein introduced to encompass phyloproteomics approaches with other identified and unidentified biomolecular features obtained by quantitative proteomics, metabolomics, lipidomics, glycoproteomics and other omics techniques. Quantphylo-based phylogeny was mapped to phylogenies inferred by concatenated ssrRNA, gGAPDH and HSP70 gene sequences to determine levels of their levels of correlation. We show the strong discriminatory power of three MS based quantitative features to accurately assign *T. cruzi* strains into their DTUs, and the suitability of this technique to reconstruct the evolutionary relationships among the DTUs of *T. cruzi*, and between this species and closely related trypanosomes. Our findings support a close congruence of the evolutionary relationships among all species and DTUs using phyloquant and sequence-based phylogenetic studies. Putative *T. cruzi* strain and DTU, and species-specific proteins at the exponential growth stage of the epimastigote life stage and multi-gene protein family expression along the parasites’ divergence are also explored.

## Methods

### Trypanosome culture

Two representative strains for the six *T. cruzi* DTUs (TcI-TcVI), one from Tcbat (TcVII), *T. cruzi marinkellei, Trypanosoma dionisii, Trypanosoma erneyi* and *Trypanosoma rangeli* (**Table 1**) were comparatively analyzed in this study. These parasites have been previously characterized using genetic markers (53, 54). The epimastigotes were grown in liver infusion tryptose (LIT) medium supplemented with 10% fetal calf serum (FCS) at 28 °C. The cells were harvested at the exponential growth phase by centrifugation at 3,000 x g for 10 min and washed twice with ice cold PBS (137 mM NaCl, 2.7 mM KCl, 10 mM Na_2_HPO_4_, 1.8 mM KH_2_PO_4_, pH 7.4). Three biological replicates of epimastigote cultures were acquired for each trypanosome to guarantee the reproducibility of the results.

**Table 1.** Summary of the *T. cruzi* strains, their discrete typing units (DTUs) and other trypanosome species analyzed in this study.

| TCC | DTU | Trypanosome strain/<br>species | Host | Locality |
| --- | --- | --- | --- | --- |
| 2158 | TcI | *T. cruzi* Sylvio X10/4 | *Homo sapiens* | Belem, PA, BR |
| 874 | TcI | *T. cruzi* G. mucurae | *Didelphis marsupialis* | Manaus, AM, BR |
| 1320 | TcII | *T. cruzi* Y | *Homo sapiens* | Rio Grande do Sul, RS, BR |
| 2188 | TcII | *T. cruzi* Esmeraldo | *Homo sapiens* | São Felipe, Bahia, BR |
| 2123 | TcIII | *T. cruzi* M-6241 cl6 | *Homo sapiens* | Belem, PA, BR |
| 844 | TcIII | *T. cruzi* | *Homo sapiens* | Carauari, AM, BR |
| 2124 | TcIV | *T. cruzi* CANIII cl1 | *Homo sapiens* | Belem, PA, BR |
| 85 | TcIV | *T. cruzi* Jose Julio | *Homo sapiens* | Amazonia, AM, BR |
| 1561 | TcV | *T. cruzi* NR cl3 | *Homo sapiens* | Bolívia |
| 2121 | TcV | *T. cruzi* 92:80 cl2 | *Homo sapiens* | Bolivia, Santa Cruz |
| 476 | TcVI | *T. cruzi* cl14 | *Triatoma infestans* | Rio Grande do Sul, BR |
| 2189 | TcVI | *T. cruzi* Cl Brener cl1 | *Triatoma infestans* | Rio Grande do Sul, BR |
| 1994 | Tcbat | *T. cruzi* bat cl1.1 | *Myotis levis* | São Paulo, SP, BR |
| 344 |  | *T. c. marinkellei* | *Carollia persipicillata* | Monte Negro, RO, BR |
| 211 |  | *T. dionisii* | *Epitesicus brasiliensis* | São Paulo, SP, BR |
| 1946 |  | *T. erneyi* | *Mops condylurus* | Moçambique, Chupanga, Sofala |
| 86 |  | *T rangeli* AM-10 | *Homo sapiens* | Amazonia, AM, BR |

### Protein extraction and digestion

The cell pellets were resuspended in protein extraction buffer containing 8 M urea and 1x protease inhibitor cocktail. The cells were lysed by three cycles of probe tip sonication on ice for 15 secs with 15 sec cooling intervals. The lysates were cleared by centrifugation at 4 °C at 18000 x g for 10 minutes, and the extracted proteins quantified by Quibit fluorometric detection method. Ammonium bicarbonate was added to a final concentration of 50 mM, followed by reduction using dithiothreitol (DTT) at a final concentration of 10 mM. The reactions were incubated for 45 minutes at 30 °C, and alkylated by the addition of iodoacetamide (IAA) to a final concentration of 40 mM and incubation for 30 min in the dark at room temperature. DTT at a final concentration of 5 mM was added to quench the reaction, and the urea diluted 10 times by the addition of NH_4_HCO_3_. Porcin trypsin was used to digestion of the proteins at a ratio of 1:50 (μg trypsin/μg protein), and the mixture incubated overnight at 30 °C. The resultant peptide mixtures were desalted with an in-house C18 micro-column of hydrophilic-lipophilic-balanced solid phase extraction. The peptides were eluted with 100 μL of 50% (v/v) acetonitrile and 0.1% (v/v) trifluoroacetic acid (TFA).

### Nano LC-MS/MS analysis

LC-MS/MS analysis were performed on an EASY-Spray PepMap^®^ 50 cm × 75 um C18 column using an Easy nLC1000 nanoflow system coupled to Orbitrap Fusion Lumos mass spectrometer (Thermo Fischer Scientific, Waltham, MA, USA). The HPLC gradient was 5 – 25% solvent B (A = 0.1% formic acid; B = 100% ACN, 0.1% formic acid) in 90 min at a flow of 300 nL/min. The most intense precursors selected from the FT MS1 full scan (resolution 120,000 full width at half-maximum (FWHM) @ m/z 200) were quadrupole-isolated and fragmented by collision-induced dissociation (CID) and detected in the dual-pressure linear ion trap with 30 as normalized collision energy. The MS1 scan range was between 380–1500 m/z, the ion count target was set to 2 × 10^5^, the MS2 ion count target was set to 1 × 10^4^, and the max injection time was 50 ms and 35 ms for MS1 and MS2, respectively. The dynamic exclusion duration was set to 10 s with a 10-ppm tolerance around the selected precursor and its isotopes. The maximum total cycle time was confined to 3s (23). All raw data have been submitted to PRIDE archive (https://www.ebi.ac.uk/pride/archive/) PXD017228.

### Peptide and Protein identification and quantification

LC-MS/MS raw files were analyzed using MaxQuant v1.5.2.8 for identification and label-free quantification (LFQ) of proteins and peptides. Using the following parameters, MS/MS spectra were searched against the combined Uniprot *T. cruzi* with 49,985 entries (downloaded, 21, July, 2017) and common contaminants protein database with a mass tolerance level of 4.5 ppm for MS and 0.5 Da for MS/MS. Enzyme specificity was set to trypsin with a maximum of two missed cleavages. Carbamidomethylation of cysteine (57.0215 Da) was set as a fixed modification, and oxidation of methionine (15.9949 Da), deamidation NQ (+ 0.9840 Da) and protein N-terminal acetylation (42.0105 Da) were selected as variable modifications. The minimum peptide length was set to 7 amino acid residues. The ‘match between runs’ feature in MaxQuant which enables peptide identifications between samples based on their accurate mass and retention time was applied with a match time window of 0.7 min and an alignment time window of 20 min. All identifications were filtered in order to achieve a protein false discovery rate (FDR) less than 1% (23, 55). Proteins identified in the reverse database, contaminants and proteins identified only by site were excluded prior to performing statistical analysis. The MS and MS/MS features considered to infer evolutionary relationships were: MS1 (quantitative values of unidentified MS1 precursor ions), iBAQ (quantitative protein expression calculated as the sum of all peptide peak intensities divided by the number of theoretically observable tryptic peptides (56), and LFQ (label free quantification based on extracted ion chromatogram area of peptides further assembled into proteins).

### Evolutionary resolution of *Schizotrypanum* lineage

Protein expression profiles evidenced by bottom up large scale mass spectrometry-based quantitative proteomic features were analyzed to discriminate *T. cruzi* DTUs and closely related trypanosomes, and to infer the evolutionary relationships of the trypanosomes in the subgenus *Shizotrypanum. T. rangeli* was included as the outgroup to *Schizotrypanum*. The m/z values of the parent-ions (MS1) are unidentified features which are independent of database search, peptide identification and genome information. The MS1 intensities in the matched features file were filtered to include all peptide intensities with a charge state of 2, 3 and 4. The cut off of charge state 1 served to differentiate the majority of peptide ions from non-peptide ions which are single-charged. The top 10 % highest MS1 intensity values which were chosen for further analysis were normalized based on the total MS1 intensities for each replicate and processed further in the Perseus computational platform (57). The m/z peptide ion intensities were transformed to log2, and filtered based on 3 valid values in at least one replicate. The Non-Assigned Numbers (NaN) were imputed from normal distribution using a down-shift of 1.8 and distribution width of 0.3. To visualize the variability of the MS1 intensities, principal component analysis (PCA) was performed on the log2 transformed MS1 intensities. To infer evolutionary relationships between the trypanosome species, Z score values (standard deviations from means) of the MS1 intensity values were calculated in Perseus, and the matrix transposed. Subsequently, decimal fractions were rounded off to the nearest integers and each negative integer substituted by a corresponding letter. Finally, Tree Analyses using New Technology (TNT) version 1.1 (58) was employed to infer evolutionary relationships using maximum parsimony. Branch statistical supports were obtained as implemented in TNT using 1000 replicates.

The second quantitative measure used in this study was the intensity Based Absolute Quantification (iBAQ). This is a measure of absolute protein amounts calculated as the sum of all peptide peak intensities divided by the number of theoretically observable tryptic peptides (56). The iBAQ values generated by MaxQuant software were normalized based on the total intensities for each replicate, loaded onto Perseus, and PCA used to visualize the variability of the proteins based on iBAQ intensities. Statistically significant iBAQ intensities were determined by analysis of variance (ANOVA) with Benjamini-Hochberg-based false discovery rate correction at an FDR of 5% (0.05). To infer evolutionary relationships based on total and statistically significant iBAQ values, maximum parsimony as implemented in TNT was used as previously described. Evolutionary trees were constructed based on total and statistically significant iBAQ values.

Thirdly, proteome-wide quantitative proteomics based on label free quantification (LFQ) of fragmented ions (MS2) were evaluated to determine the discriminatory power of the total and significantly expressed proteins between the strains and species, and to reconstruct the evolutionary relationships of the trypanosomes. The LFQ values were analyzed by Perseus as previously described. To determine proteins with significant changes in abundances, analysis of variance (ANOVA) was applied with Benjamini-Hochberg-based false discovery rate correction at an FDR of 5% (0.05). The total and significantly expressed protein LFQs were normalized using the Z score and the evolutionary relationships inferred using parsimony as previously described.

### Phylogenetic clustering

Three concatenated genes; glycosomal glyceraldehyde 3-phosphate dehydrogenase (gGAPDH), v7v8 variable region of the SSU rRNA and HSP70 were selected to construct a phylogenetic reference tree. These gene sequences have been routinely used to discriminate and infer phylogenetic relationships among trypanosomes of *Schizotrypanum* and *T. rangeli* (53). Orthologs of the three genes were obtained from the available genomes (complete and drafts) using a standalone BLASTn (v2.2.31+). The genomes from *T. cruzi* (Sylvio X10), *T. cruzi* (CL Brener) and *T. cruzi* (Esmeraldo) are available in TryTrypDB (59). *T. cruzi* (Y) genome is available in NCBI-Genenank (60). The genome drafts from *T. dionisii* (TCC211), *T. erneyi* (TCC1946), *T. rangeli* AM80 (TCC86), *T. cruzi marinkellei* (TCC344), *T. cruzi G* (TCC874) and *T. cruzi* M-6241 (TCC2123) have been generated in our laboratories within the ATOL (Assembly of Life, NSF-USA) and TCC-USP (Brazil) projects and were obtained as previously described (61). Genes for *T. cruzi* MT3869 (TCC844), *T. cruzi* CANIII (TCC2124), *T. cruzi* Jose Julio (TCC85), *T. cruzi* 92:80 (TCC2121) were retrieved from NCBI repository using BLASTn.

SSU rRNA gene from *T. cruzi* Cl *Brener* (TCC2189), gGAPDH genes from *T. cruzi* NR Cl1 (TCC1561) and *T. cruzi* Cl14 (TCC476), and HSP70 genes from *T. cruzi* 92:80 (TCC2121), *T. cruzi* NR Cl1 (TCC1561) and *T. cruzi* Cl14 (TCC476) were amplified by PCR as previously described (53, 62, 63), and subsequently sequenced. All gene sequences were aligned using CLUSTAL v2.1 within the Seaview v.4.5.4 graphical tool software (64). Manual refining of the alignment was performed prior to inferring phylogenies by the parsimony method using TNT (58). Branch statistical supports were obtained using bootstrap with 1000 replicates.

### Phylogenetic and phyloquant correlation

Statistical correlation between phylomics and phylogenetics were determined by mantel test using the Analysis of Phylogenetics and Evolution (ape) package of the R statistical language environment for statistical computing (65). Distance matrices based on the generated evolutionary trees were generated using Patristic Distance Matrix and statistical strength of correlation between the matrices generated from phylogenetics and phyloquant-based clustering calculated by the Mantel test (66), using the APE package implemented in the R statistical software.

### Putative strain/DTU/species-specific protein identification

Putative strain/DTU/species-specific proteins exclusively expressed in epimastigotes from exponential growth phase of the trypanosomes were determined. The proteins were considered specific if the sum of the protein intensity of all the other strains and/or DTUs was zero, while the sum of the DTU/species under consideration was more than zero considering at least two biological replicates. For this analysis, the NaN values were imputed with a zero value in the Perseus platform.

### Divergence of multigene protein family

Pearson correlation was used to evaluate degrees of correlation between protein expression and dates of divergence of the trypanosomes assayed based on LFQ-based phylogeny. Selected trypanosome multi-gene protein families were considered for this analysis, and included ribosomal proteins, trans-sialidases, ABC-transporter, gp63, heat shock proteins (HSPs), dispersed gene family (DGF-1), paraflagellar-rod proteins, cytoskeletonAssociatedProtein5.5 and retrotransposon hot spot (RHS) proteins. Correlation was performed and visualized in Hmisc and corrplot packages, respectively, implemented in R statistical software (67, 68).

## Results

### Mass spectrometry results

The evolutionary relationships of *T. cruzi* strains representative of all DTUs and closely related trypanosomes of the subgenus *Schizotrypanum* were investigated using a comprehensive bottom up mass spectrometry-based quantitative proteomics combined with statistical and computational analyses. A schematic flowchart of the experimental procedures applied in this study is summarized in **Figure 1**. All trypanosome strains/species were harvested in the epimastigote life stage collected at the exponential growth phase of the cultures. Parasite proteins were extracted, digested with trypsin and analyzed by nLC-MS/MS. The raw MS/MS files were analyzed with MaxQuant v.1.5.3.8 to identify and quantify proteins using the ANDROMEDA search engine against the Uniprot *T. cruzi* protein database using label free quantification. Proteome-wide quantitative proteomics using unidentified (MS1) and identified (iBAQ and LFQ) mass spectrometry features were evaluated to determine the evolutionary relationships of the genetically highly diverse *T. cruzi* strains and closely related trypanosome species. This approach was termed ‘*phyloquant*’ to include all studies hereafter which infer evolutionary relationships or biologically meaningful classifications based on mass spectrometry-based intensities of biomolecules including proteins, lipids, proteins, glycans, glycoproteins.

**Figure 1.**
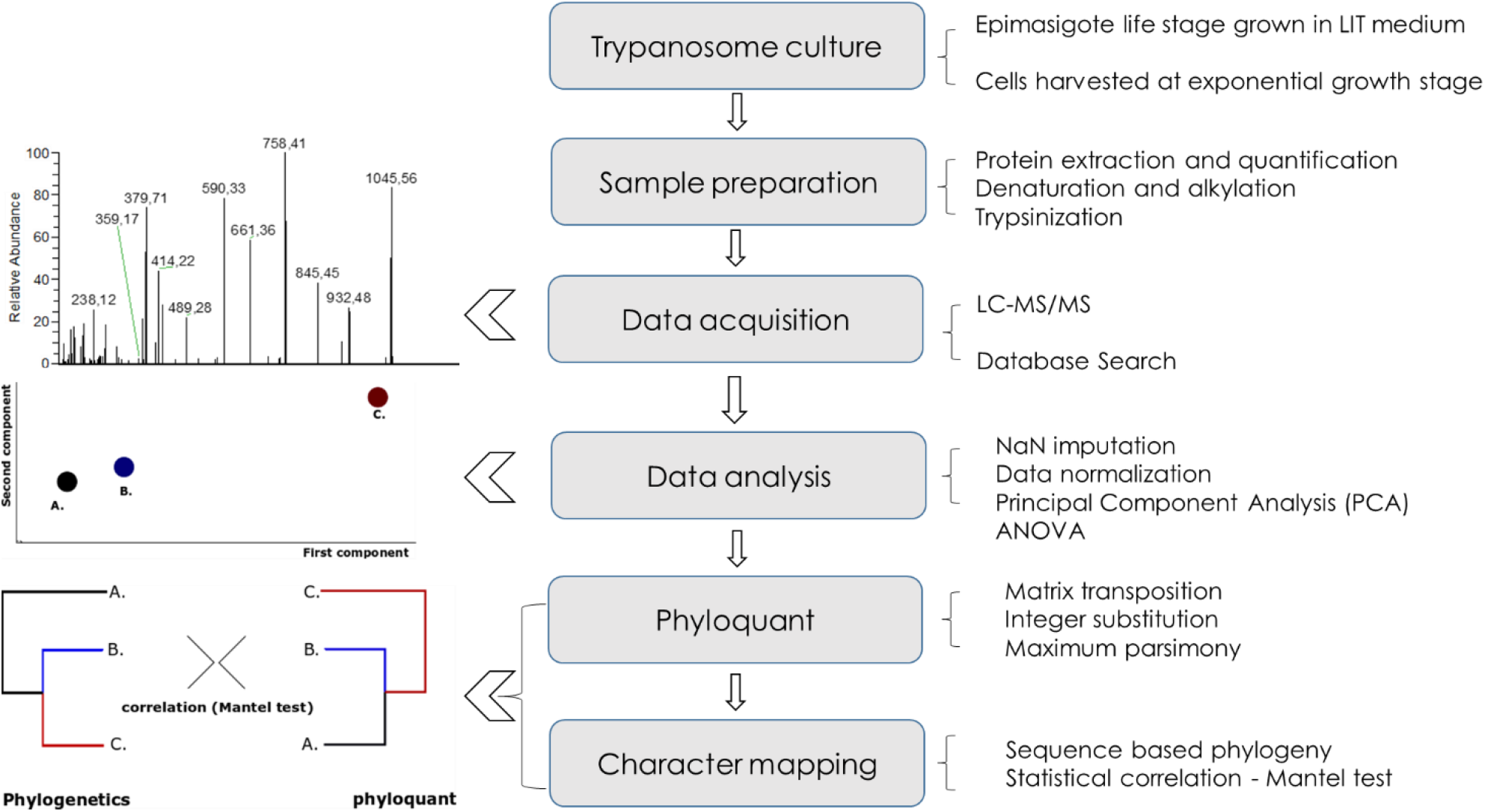
Experimental workflow for the phyloquant approach. Trypanosomes grown in complete LIT medium were harvested at the exponential proliferative phase of the epimastigote life stage. Proteins were extracted in 8M urea lysis buffer in the presence of protease inhibitors, digested with trypsin and the resultant peptides analyzed by nLC-MS/MS. Normalized (**A**) MS1, (**B**) iBAQ and (**C**) LFQ values were used to infer evolutionary relationships using the parsimony algorithm. Phylogeny based on three concatenated genes, GAPDH, V7V8 ssrDNA and HSP70 was chosen to perform character mapping of the evolutionary relationships based on proteome-wide mass spectrometry-based features. Statistical levels of correlation between phyloquant and phylogenetic approaches were inferred using the mantel test.

A total of 527,067 MS1 values from precursor ions with a charge state of ≥ 2 were quantified from the 17 trypanosomatids in all the biological replicates evaluated. Of these, 450,372 precursor ion intensities had a charge state of 2, 3, and 4 (**Supplementary Table 1**). The most intense MS1 values (10%) with a charge state of 2,3,4 were chosen for further analysis and their normalized values were subsequently analyzed in Perseus software platform. Following filtering of the data to include only intensities with 3 valid values in at least one strain/species, a total of 27,446 MS1 intensities were chosen for evolutionary inference. A total of 3,889 proteins values were identified and quantified in this study (**Supplementary Table 2**). Of these, 2224 and 1990 iBAQ and LFQ values, respectively, were statistically significant (FDR less than 0.05), with 3 valid values in at least one group. All the quantitative proteomic values from MS1, iBAQ and LFQ normalized by z-square, and parsimony, a phylogenetic algorithm, used to infer evolutionary relationships.

### Variation of protein expression among trypanosomes

A multivariate data analysis based on principal component analysis (PCA) was employed to explore the variation in protein expression between trypanosomes obtained by LC-MS/MS. Variations of Log(2) transformed MS1, iBAQ and LFQ were visualized by the first and second principal components (**Figure 2**). PCA on MS1 intensities placed the *T. cruzi* DTUs into distinct clusters separated from the bat allied *T. dionisii* and *T. erneyi*, and the generalist *T. rangeli*. This clustering was based on the first and the second principal components, which represented 9.1% and 6.3% of the total variation in the dataset, respectively (**Figure 2A**). TcIV and Tcbat tightly clustering together, the parental genotype (TcII) and the hybrid genotypes (TcV and TcVI) clustered closer together. *T. cruzi marinkellei*, a subspecies of *T. cruzi* strongly linked to bats, was placed closer to *T. cruzi* DTUs (TcI and TcIII) than to the allied bat trypanosomes *T. dionisii* and *T. erneyii*, which are the most basal species of *Schizotrypanum*, which formed distinct clusters. As expected, the generalist *T. rangeli*, included as an outgroup in this study, clustered separately from all the species of *Schizotrypanum*. Proteomic variation based on iBAQ values resulted in similar clustering (**Figure 2B**), showing separation of *T. cruzi* DTUs from the allied bat trypanosomes and *T. rangeli*. Distinct separation of *T. cruzi* DTUs by PCA showed four sub-clusters; TcI and Tcbat formed a well separated cluster based on the second principal component representing 11.7% of the total variation in the dataset, while strains representative of TcII, TcV and TcVI formed a cluster separate from strains representative of TcIII and TcIV. *T. cruzi marinkellei* was placed closer to the bat allied *T. dionisii* and *T. erneyi* than to *T. cruzi* DTUs. *T. rangeli* was placed furthest away from all *T. cruzi* DTUs. PCA analyses based on LFQ features resulted in three distinct clusters based on the first and second components representing 26.4% and 9.7% of the total variability in the dataset, respectively (**Figure 2C**). TcI and Tcbat were placed closer together, while the parental TcII and hybrid genotypes TcV and TcVI clustered together. TcIII and TcIV were separated from TcI-Tcbat and TcII-TcV/TcVI, forming separate clusteres. *T. c. marinkellei* was placed between *T. cruzi* cruzi clade and *T. dionisii, T. erneyii* and the more distant cluster of *T. rangeli*.

**Figure 2.**
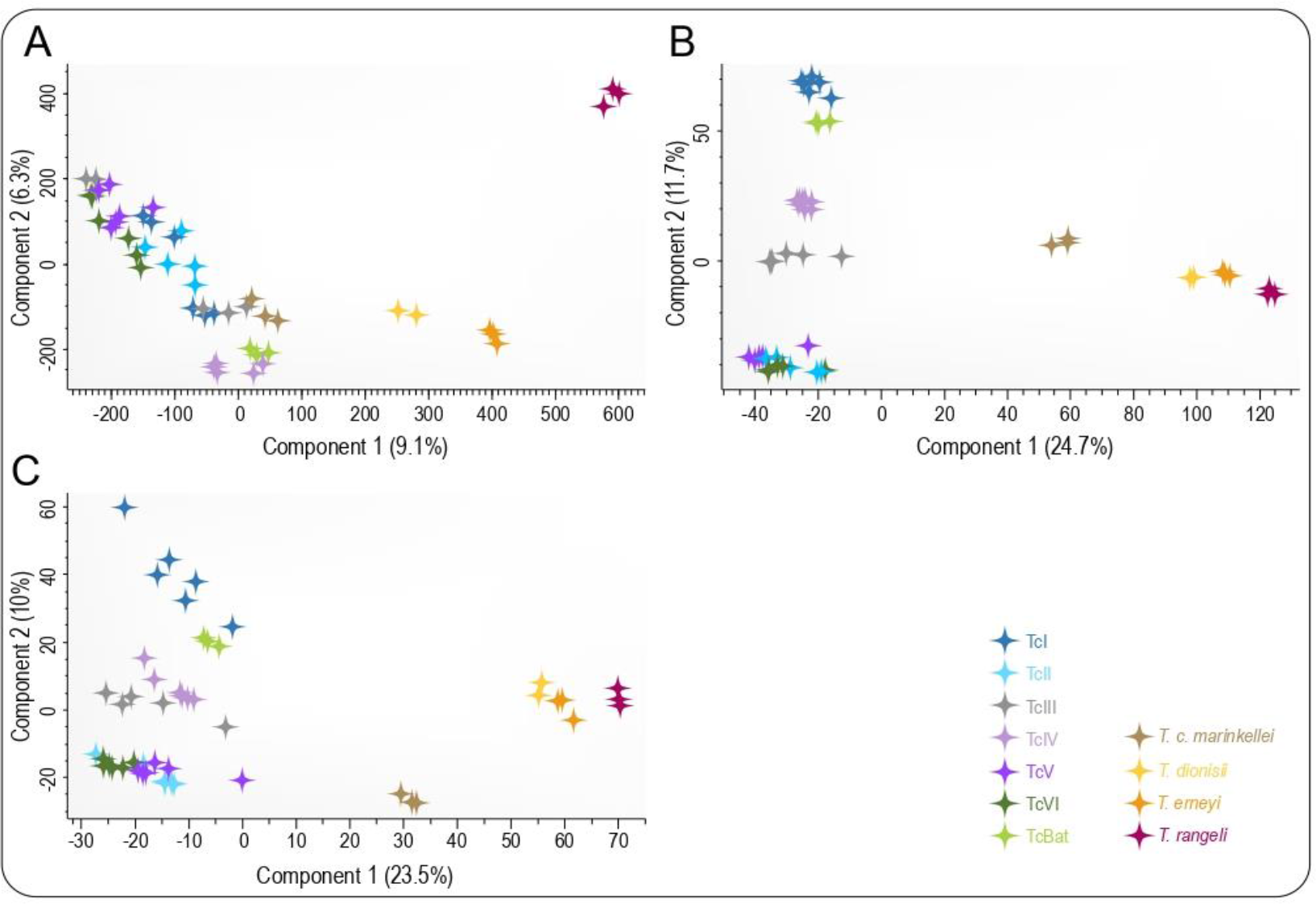
Principal component analysis of *T. cruzi* DTUs and bat allied trypanosomes; *T. c. marinkellei* (grey), *T. dionisii* (green), *T. erneyi* (black), and the generalist *T. rangeli* (red). PCA was performed on normalized **A**) MS1 values, **B**) iBAQ and **C**) LFQ proteomic features.

### Protein expression-based evolutionary inference *(‘phyloquant’)*

The term *‘phyloquant’* is described here as the method of inferring the evolutionary relationships between strains/species of trypanosomes based on quantitative features derived from mass spectrometry-based analyses. In addition, this term can be applied to similar studies which use quantitative mass spectrometry combined with a phylogenetic algorithm to infer biologically meaningful classifications of samples or evolutionary relationships of different strains/species. Contrary to traditional molecular phylogeny which determines the evolutionary distances of organisms based on DNA or predicted protein sequence information, phyloquant is a numeric taxonomy which infers evolutionary relationships based on masses and intensities of ions or fragments quantified by mass spectrometry. We propose the term *‘phyloquant’* to also encompasses both identified and unidentified mass spectrometry-based fingerprints from peptides, proteins, metabolites, lipids, glycopeptides, glycans, phosphoproteins and other omics based on quantitative mass spectrometry. In this proof of concept study, we evaluated the potential application of quantitative proteomic-based data for lineage assignment and inference of evolutionary relationships of *T. cruzi* strains and closely related trypanosomes. Phylogenies were inferred by parsimony using numeric values from the most intense unidentified precursor ions (m/z) values; and total and statistically significant intensities of identified MS-based proteomic features. Subsequently, character mapping of the phyloquant approach was performed based on phylogenies from concatenated sequences of three gene loci; gGAPDH, v7-v8 region of the ssu rRNA and HSP.

Phyloquant clustering inferred from the top 10% high intense MS1 m/z intensities showed the separation of *T. cruzi* DTUs from the closely phylogenetically related trypanosomes strongly linked to bats, *T. c. marinkellei, T. erneyi* and *T. dionisii* (**Figure 3A**). *T. c. marinkellei* formed the basal clade to *T. cruzi* DTUs, while TcI-Tcbat sister clade was placed basal to the *T. cruzi cruzi* clade. *T. cruzi* strains were grouped in three major clusters; TcI-Tcbat, TcIV, and TcII/III/V/VI. Strains representative of TcI, TcIV, TcV and TcVI clustered to form monophyletic clades, while TcII and TcIII strains were the most divergent within DTUs, forming paraphyletic clades distributed into TcV and TcVI. Tcbat was placed sister to TcI forming a clustering congruent with evolutionary relationships inferred by various genetic markers including cytb, gGAPDH, SSU rRNA, H2B and many other gene sequences (54, 69).

**Figure 3.**
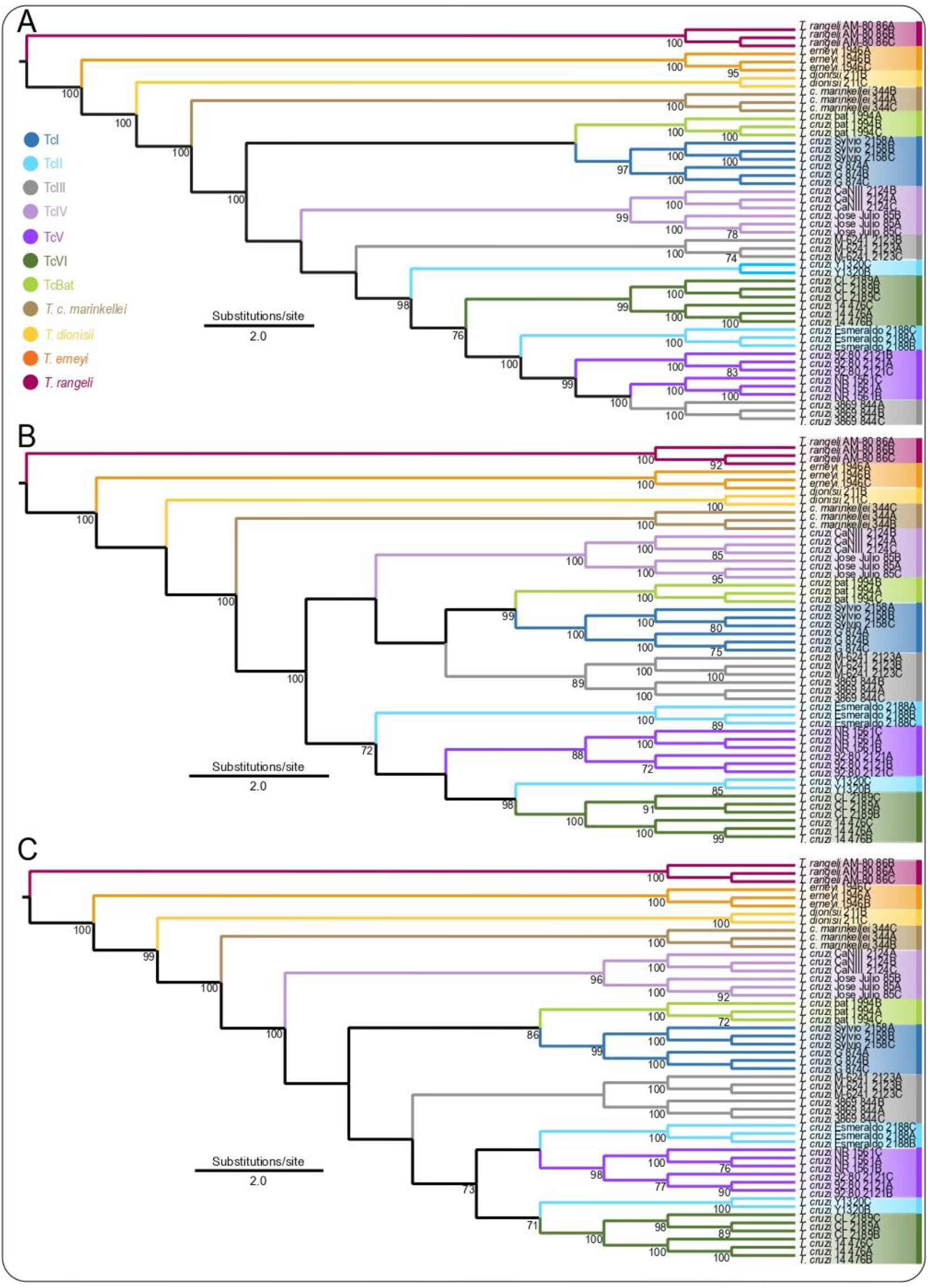
Phyloquant analysis of trypanosomes of the *Schizotrypanum* subgenus based on differentially regulated (**A**) MS1, (**B**) iBAQ and (**C**) LFQ values inferred by the parsimony evolutionary algorithm. Numbers on branches represent bootstrap support above 70 % estimated with 1000 replicates.

Phyloquant inferred from statistically significant iBAQ values using a Benjamini-Horchberg based FDR at an FDR < 0.05 showed the separation of *T. cruzi* DTUs from the closely related *T. c. marinkellei, T. dionisii* and *T. erneyi* and the outgroup *T. rangeli*, supported by high bootstrap values (**Figure 3B**). Moreover, iBAQ based phylomics demonstrated a high inter DTU discriminatory power, enabling the assignment of *T. cruzi* strains to their monophyletic DTU clades, with the exception of DTU TcII, which remained polyphyletic. *T. c. marinkellei* formed the basal clade to *T. cruzi* DTUs, supported by a high bootstrap value (100%). *T. cruzi* DTUs nested into two major clusters; TcIV/TcI-Tcbat/TcIII and TcII/TcV/TcVI. TcIV formed the basal clade to the first cluster, supported by a high bootstrap value (100%). TcI-Tcbat sister group was more placed sister to TcIII. A clustering congruent to MS1 based phylomics showed *T. cruzi Esmeraldo* and *T. cruzi* Y strains, (TCC2188) were more related to TcV and TcVI, respectively. This clustering was conserved in phyoquant analysis based on total iBAQ values, albeit with different bootstrap support values (**Supplementary figure 2A**).

Phylomics based on statistically significant LFQ values also showed that *T. cruzi* DTUs were nested together, well separated from the bat allied trypanosome species and the generalist *T. rangeli* with high support values (**Figure 3C**). Within *T. cruzi cruzi* clade, 10 of the 12 *T. cruzi* strains were assigned to their monophyletic DTU clades, with the exception of strains representative of TcII which were nested in polyphyletic clusters. Three main clusters of *T. cruzi cruzi* clade were formed based on statistically significant LFQ values; TcIV strains formed the basal clade to *T. cruzi* DTUs, while TcI_Tcbat strains formed a sister group, corroborated by MS1 and iBAQ-based phylomics. The parental and hybbrid genetypes formed the third clade haboring monophyletic TcIII, TcV and TcVI DTUs, and the polyphyletic TcII. *T. cruzi* esmeraldo was placed closer to TcV, while *T. cruzi Y* strain showed a close relationship with TcVI. Phylomics based on total LFQ values showed similar clustering patterns to differentially regulated LFQ values, however TcIV and TcIII were not clustered as monophyletic clades (**Supplementary Figure 2B**).

### Phyloquant shows high correlation with phylogenetics

To infer evolutionary correlation between phyloquant and phylogenetics, one random replicate for each trypanosome strain/species was used to construct evolutionary trees based on MS1, iBAQ and LFQ values. Subsequently, generated phylogeny trees were mapped to a molecular phylogeny inferred by parsimony based on concatenated gGAPDH, ssrRNA and HSP70 genes (**Supplementary Figure 1**). Phylogenies inferred by the concatenated gene sequences have previously been employed to infer phylogenetic relationships aimed at resolving relationships between *Schizotrypanum* lineage and the Tra [Tve-Tco] lineage (53) comprising *T. rangeli, T. conorhini, T. vespertilionis* and other Old World trypanosomes.

Levels of correlation were statistically evaluated using the non-parametric mantel test using analyses of phylogenetics (APE) and implemented in R statistical software. Mantel test gave a statistically significant correlation (p-value of 0.0009 at an alpha of 0.05), showing close correlation between three evolutionary trees based on MS1, iBAQ and LFQ and phylogenetics. This was determined based on 1000 permutations. The statistically significant correlation between phylogenies derived from protein expression levels and molecular based phylogenies illustrate the high correlation between *T. cruzi* protein expression and infraspecific genetic subdivisions, as earlier illustrated by Telleria *et. al*. (49).

### DTU and species-specific proteins

The LFQ values were separately summed for each trypanosome species or *T. cruzi* DTU, and the sum of the remaining species and DTUs were summed separately. A protein was considered unique if it was identified exclusively in one DTU or species and the sum of the remaining species was 0. The identification of proteins unique to each species, DTU or strain does not imply the absence of the protein, but is related to the abundance below the limit of analytical detection, the presence of modified peptides and/or unique peptide sequences. The putative *T. cruzi* specific strain/DTU and species-specific proteins identified in this study is summarized in **Table 2**.

**Table 2.** Putative strain/DTU and species-specific proteins identified at the exponential growth phase (epimastigote forms) of *T. cruzi* DTUs and closely related trypanosome species

| Strain/species | DTU | Strain/species<br>specific protein | DTU specific protein |
| --- | --- | --- | --- |
| Tcc2158_*T. cruzi* Sylvio X10/4 | TcI | 6 |  |
| TCC874_*T. cruzi* G | TcI | 10 | 42 |
| TCC2188_*T. cruzi* esmeraldo | TcII | 17 |  |
| TCC1320_*T. cruzi* Y | TcII | 1 | 1 |
| TCC2123_*T. cruzi* M-6241 | TcIII | 1 |  |
| TCC844_*T. cruzi* 3869 | TcIII | 4 | 6 |
| TCC2124_*T. cruzi* CANIII | TcIV | 5 |  |
| TCC85_*T. cruzi* Jose Julio | TcIV |  | 3 |
| TCC2121_*T. cruzi* 92:80 | TcV | 1 |  |
| TCC1561_*T. cruzi* NR Cl3 | TcV | 2 | 3 |
| TCC2189_*T. cruzi* CL Brener | TcVI | 1 |  |
| TCC476_*T. cruzi* Cl14 | TcVI | 2 | 3 |
| TCC1994_Tcbat | Tcbat | 17 |  |
| TCC344_*T. c. marinkellei* |  | 279 |  |
| TCC211_*T. dionisii* |  | 8 |  |
| TCC1946_*T. erneyi* |  | 8 |  |
| TCC86_*T. rangeli* |  | 4 |  |

A total of 42 proteins were exclusively identified in TcI strains represented by *T. cruzi* Silvio and *T. cruzi* G quantified in at least two replicates for each strain. Multicopy gene families were among the uniquely identified proteins specific to TcI, including ABC transporter (V5D8T8; TCDM_08166) and 2 Retrotransposon hotspot (RHS) proteins (V5BCL4; V5AKS1). Cathepsin B-like protease (O61066; tccb) was uniquely identified in TcI, with 4 unique peptides out of 7 peptides identified, assigned to two proteins (O61066; Q95VX8) sharing 99.52% sequence similarity. Cathepsin B-like protease is a single copy gene expressed in all the life stages of *T. cruzi* (70), and is involved in intracellular protein catabolism (71). Lipophosphoglycan biosynthetic protein (V5BKG0; TCDM_03038) was identified with 6 unique peptides from a total of 35 identified peptides, which were assigned to one protein accession number involved in protein folding. Importin subunit alpha (V5B924; TCDM_01484) specific to TcI was identified, with 5 unique peptides from a total of 11 identified peptides assigned to one protein accession number. This protein has a nuclear import signal receptor involved in the import of proteins into the nucleus. Moreover, 6 and 10 putative *T. cruzi* sylvio and *T. cruzi* G strain specific proteins, respectively, were identified. Among the *T. cruzi* G strain specific proteins were included two RHS proteins (V5BNA2; TCDM_02054 and V5B9S5; TCDM_11312) and proteasome beta 3 subunit (V5BGL9; TCDM_14499). Putative *T. cruzi* Sylvio strain specific proteins included Retrotransposon hot spot (RHS) protein, putative (Q4DQ78; Tc00.1047053510543.90) and Chaperonin alpha subunit (V5B209; TCDM_04101).

A total of 18 proteins were exclusively identified in *T. cruzi* esmeraldo which were quantified in at least two replicates. Of these, one protein was identified in one replicate in *T. cruzi* Y and in three replicates for *T. cruzi* esmeraldo, and was considered a putative TcII specific protein. This protein was not characterized (Q4D7I8, Tc00.1047053509961.30), and GO terms using UniProt showed that it’s involvement in tRNA splicing via endonucleolytic cleavage and ligation. This protein was identified with 4 peptides, all unique, and were assigned to 15 protein groups. Three of 18 *T. cruzi* esmeraldo-specific proteins had functions involving oxidoreductase activity, important in the maintenance of redox status. These included Glutamate dehydrogenase (Q4DWV8; Tc00.1047053507875.20), Thiol-dependent reductase 1 (G0Z1H4) (Q2TJB3) and Prostaglandin F2alpha synthase (G0Z1C2).

A total of 6 proteins were identified exclusively expressed in TcIII strains. Dihydrolipoamide acetyltransferase putative was identified in all six triplicates of TcIII, with two unique peptides from a total of 8 identified peptides, which were assigned to one protein accession number (Q4DYI5, Tc00.1047053511367.70). This protein has an acetyl transferase activity. Leucine amino peptidase (fragment) (H2CS44, LAP) was identified uniquely for *T. cruzi* M-6241 in all three triplicates with 1 unique peptide from 6 identified peptides assigned to 2 protein accession numbers (H2CS44, A8W0Z4) which shared 100% sequence identity. Four putative strain specific proteins identified in all three triplicates for *T. cruzi* 3869 included a retrotransposon hotspot (RHS) protein (V5AQH5, TCDM_12607), an uncharacterized protein (Q4E183; Tc00.1047053506459.270), a Serine palmitoyltransferase, putative (Q4DID6; Tc00.1047053506959.70) and Cyclophilin (V5B9Y6; TCDM_09432). Cyclophilin protein was assigned to 4 protein accession numbers; V5B9Y6 (TCDM_09432), Q4DJ19 (Tc00.1047053504797.100), Q6V7K8 and Q4DM35 (Tc00.1047053511577.40), which shared high sequence homology > 98.05%. Cyclophylin is a peptidyl-prolyl cis/trans isomerase enzyme with diverse biological functions, including protein folding, stress response, immune modulation, regulation of parasite cell death, and signal transduction. In *T. cruzi*, cyclophilin gene family of 15 paralogous genes has been reported (72). The upregulation of a *T. cruzi* cyclophilin, TcCyP19, was demonstrated in benznidazole resistant populations (73, 74). Indeed, TcIII (represented by strain 893) is among the DTUs were resistance to benznidazole has been reported (75). The anti-parasitic activity of Cyclosporin A (CsA), an immunosuppressive drug which targets cyclophilins, has been demonstrated in *T. cruzi* (76, 77).

Three TcIV-putative proteins were uniquely identified with at least 5 valid values in total. These included RHS protein (V5B362; TCDM_12365), an uncharacterized protein (K2NRV8; MOQ_002111) and a Glusose-6-phosphate isomerase (Q6RZZ5; Gpi). Five putative strain specific proteins were identified for *T. cruzi* CANIII with at least 2 valid values. These included two RHS proteins (V5CIH1, Q4D2I5), an uncharacterized protein (I3QFA4), clathrin light chain (Q4DYJ4) and mevalonate kinase (I3QEA0).

Three putative TcV specific proteins were identified with at least 5 valid values. The three proteins were from the multigene family of RHS proteins; Q4DBD1(Tc00.1047053504099.10), Q4CKU7 (Tc00.1047053399997.10), Q4CU66 (Tc00.1047053463155.20). In addition, two putative RHS proteins (Q4D0T5; Tc00.1047053506845.60 and Q4CL80; Tc00.1047053446325.9) were separately identified in at least two replicates for *T. cruzi* NR and *T. cruzi* 92:80 strains, respectively. A surface protein-2 (Q86DL6) with an exo-alpha sialidase activity was identified with in three *T. cruzi* 92:80 replicates with 6 peptides, of which 2 were unique and assigned to 2 proteins (Q86DL6; Q86DL7) with a 100% shared homology based on sequence coverage. This protein is involved in pathogenesis of the parasite. Strain specific proteins for *T. cruzi* NR and *T. cruzi* 92:80 identified only in each strain in at least two replicates included an uncharacterized protein (V5DS84; TCDM_01067) and a putative trans-sialidase (K2MKD3; MOQ_010145), respectively.

One protein was identified unique to TcVI with at least three valid values for each strain. This was an uncharacterized protein (Q4E501; Tc00.1047053508873.420). Putative strain specific protein for *T. cruzi* Cl14 with at least two valid values included a trans-sialidase, putative (Q4D8B3; Tc00.1047053505919.20) and a Multidrug resistance protein E (V5BK30; TCDM_04857). One putative *T. cruzi* ClBrener specific protein was identified in our analysis; an uncharacterized protein (K2MR20; MOQ_008196) with a GO annotation as an integral component of membrane.

We identified 17 putative Tcbat proteins considering at least two valid values out of three replicates of the strain Tcbat. A trypanothione reductase (A0A0M3YE01) was identified with one unique peptide from 23 identified peptides assigned to one protein. This protein is involved in the maintenance of the cell redox environment, and has been proposed as a putative drug target against Chagas disease (78). ABCG1 protein (A0A0C5C2U8) was identified unique to Tcbat with one unique peptide from a total of 16 identified peptides assigned to three protein accession numbers (A0A0C5C2U8, A0A0C5BZ95, A0A0C5BV51), which shared > 99.7% sequence similarity. ATP-binding cassette (ABC) transporters use the hydrolysis of ATP to translocate compounds across biological membranes. Over expression of TcABCG1 has been implicated in the resistance of *T. cruzi* to benznidazole (75, 79). A trans-sialidase (Q4D095) and two surface proteins; (Q6WAZ7, sp1) and 82-kDa surface antigen (A0A0A7DWF9) with exo-alpha sialidase activity involved in pathogenesis, were among the proteins identified exclusive to Tcbat. A study by Maeda and colleagues (80) reported the surface molecules and infectivity of Tcbat metacyclic trypomastigotes to human HELA cells and mice models. In this study, Tcbat was shown to be infective to HELA cells, but its infectivity to mice models was lower compared to CL and G strains. Gp82, involved in host cell invasion by the induction of lysosomal exocytosis, was shown to be expressed and highly conserved in Tcbat, CL and G strains (80). A study by Marcili and group also showed the infectivity of Tcbat in human cells and in experimentally infected mice (31), albeit with low parasitaemia. A mixed infection of Tcbat and TcI has also been reported in a child from northwestern Colombia (81), and Tcbat has also been demonstrated in mummies from Colombia desert by molecular paleoepidemiological tools (82). The identification of putative Tcbat proteins or protein expression profile can help to understand this atypical DTU of *T. cruzi*, to date never isolated from naturally infected humans, and to study of the Tcbat epidemiology. A total of 279 proteins were uniquely identified in *T. c. marinkellei* in at least two replicates. These proteins play diverse biological functions that include roles in nucleoside metabolic process, protein folding, cysteine biosynthetic process from serine, cell redox homeostasis and arginine biosynthetic process. Interestingly, superoxide dismutase (K2MYD1; MOQ_008923) was among the proteins identified uniquely expressed in *T. c. marinkellei* based on a 2DIGE coupled to MS/MS study by Telleria *et al*. (49). In our analysis, superoxide dismutase was identified with 4 unique peptides out of 10 identified peptides, which were assigned to two proteins (A0A0M3YIG0; K2MYD1) sharing 99.52% identity. Two structural proteins related to the flagellum, the Paraflagellar rod protein 3, putative (K2LTU6; MOQ_010227) and Paraflagellar rod component, putative (K2M6S7), were identified unique to *T. c. marinkellei*. Paraflagellar rod protein 3 was identified with 52 identified peptides, of which 2 were unique. Other *T. c. marinkellei* putative specific proteins identified in this study included elongation factor 2, putative (K2MQL8, MOQ_006823), Glucose-regulated protein 78, putative (K2NLB6; MOQ_006525) and ATP-dependent Clp protease subunit, heat shock protein 100 (HSP100), putative (K2MHR5, MOQ_009649), among others (**Supplementary Table 3**).

Eight proteins were uniquely assigned to *T. dionisii* with identification and quantification in the two replicates assayed. These included Cathepsin L-like protein (J3JZZ3), Trypanothione reductase (Q963S3), RHS protein, putative (Q4E2L4) and Microtubule-associated protein (Q9U9R2).

Eight (8) putative *T. erneyi* proteins were identified exclusively expressed in *T. erneyi* including cytosolic Glyceraldehyde 3-phosphate dehydrogenase, putative (Q4CLF1, Tc00.1047053508537.10), Acetonitate hydratase (K2MVY5, MOQ_004887), NAD synthase, putative (K2N5H6, MOQ_002882), a transialidase (Q9BHJ5, TCTS), Nicotinate phosphoribosynthase, putative (K2N3L8, MOQ_003643), Choline/carnitine O-acetyltransferase (V5BQH8), Pumilio/PUF RNA binding protein 2 (V5BJ54) and an uncharacterized protein (V5BB64, TCDM_06850).

Four proteins were uniquely identified and quantified in *T. rangeli*, detected in all the three replicates. A Heat-shock protein 70 (V5JFQ1/V5JFY9) was identified with 3 unique peptides from a total of 34 identified peptides. These peptides were assigned to two protein accession numbers sharing 98.91% identity. Another protein exclusively identified in *T. rangeli* was Cathepsin L-like protein (J3JZZ4; CatL-like) with 7 unique peptides. Cathepsin L-like protein is a cysteine protease which, in protozoan parasites including trypanosomes, have been reported to play key roles in multiplication and cell differentiation, metabolism, and virulence, including involvement in invasion, evasion and modulation of host immune responses (83–85). This gene occurs in multiple copies (86), and has previously been proposed in the diagnosis, typing and evolutionary studies of *T. rangeli* (87). In addition, this enzyme is a validated drug target and is a vaccine candidate for *T. cruzi* (88). The 60S acidic ribosomal protein P2 (R9TL46), identified with 5 unique peptides from the 6 peptides quantified, was assigned to 6 protein groups (Q66VD2, E9BMC1; A4I600; A4I5Z9, O43940) which shared a sequence similarity greater than 99%, with the exception of R9TL46 with an identity > 71%. This protein is a structural constituent of ribosomes and plays a role in translational elongation. Glycosomal glyceraldehyde phosphate dehydrogenase (C4PLA3, gGAPDH) was identified with two unique peptides out of 13 identified peptides assigned to three proteins (C4PLA3, C4PLA2, A0A165D005). BLASTp analyses showed a high sequence similarity (>91%) between the three proteins. gGAPDH (C4PLA3) was identified as exclusively expressed protein in *T. rangeli*. Because gGAPDH is a very abundant, well conserved protein that is constitutively expressed in all trypanosomatids, the identification of this protein in *T. rangeli* can be associated to unique peptide sequence. These putative specific peptide sequences offer potential to be evaluated as *T. rangeli* specific epitopes in the screening of mixed infections.

### Divergence

Correlation analysis of protein expression and divergence of T. cruzi and allied trypanosome species was analyzed by Pearson correlation. In this analysis, *T. rangeli* was used as the most common recent ancestor to the *Schizotrypanum* clade. Proteins such as trans-sialidases, retrotransposon hot spot (RHSP) proteins and ABC transporter proteins showed an increase in expression along the divergence of the parasites from the common ancestor, at a p-value < 0.05. On the other hand, heat shock proteins, ribosomal protein, kinases and phosphatases showed proteins which were positively and negatively correlated to divergence (**Supplementary figure 3–6**).

## Discussion

In this study, a novel approach to classify and infer evolutionary relationships between trypanosomatids of the Schizotrypanum subgenus is described. This approach is based on different biomolecular features (MS1, iBAQ and LFQ) obtained by bottom up large scale mass spectrometry-based quantitative proteomics analysis followed by parsimony, an evolutionary algorithm to infer evolutionary distances between strains/species. This new analysis requires a new nomenclature that describes evolutionary inference of different species or strains using different features based on mass spectrometry. We propose the term “phyloquant” to infer methodologies which use biomolecular features obtained by mass spectrometry-based methods to infer evolutionary relationships between organisms. Phyloquant encompasses proteomics, metabolomics, lipidomics, glycomis and other omics techniques based on mass spectrometry. Historically, evolutionary relationships are deduced from DNA/RNA or predicted protein sequence data based respectively on nucleotide and amino acid sequence alignments from different organisms for their comparisons. Evolutionary trees build using phylogenetic approaches use neighbour joining (NJ) (89), maximum parsimony (MP) (90), maximum likelihood (ML) (91), and Bayesian inference methods (92).

The term phyloproteomic has been used to compare biological clustering using phylogenetic algorithms to protein expression based on mass spectrometry-based data and on protein sequence-based clustering. Abu-Asad and colleagues used the word phyloproteomics to classify serum cancer and non-cancerous samples using parsing and a phylogenetic algorithm (93). In particular, SELDI-TOF MS data were obtained from serum collected from cancer and non-cancer patients and the m/z values were sorted for polarity in order to identify derived vs ancestral peaks using MIX, a phylogenetic algorithm. Another study investigated *C. jejuni* ssp. *jejuni* to develop a phyloproteomic typing scheme that combines the analysis of variable masses observed during intact cell MALDI-TOF MS with ribosomal MLST and whole genome MLST database-deduced isoform lists (94). Another study used the term phyloproteomics to infer evolutionary relationships of bacteria based on mass spectrometry-based identification and protein sequence information (95). The term phyloquant proposed in the present study unifies any method that uses quantitative omics data derived from mass spectrometry. It is important to notice that the correspondence between any phyloquant approach and phylogenetic evolutionary relationships should be evaluated in each organism under study, and differences or similarities arising from this comparison holds important biological meanings.

In this study, we tested the validity of different biomolecular features obtained by LC-MS/MS bottom up quantitative proteomic strategy to explore the evolutionary diversity of trypanosomes nested into the subgenus *Schyzotrypanum* clade and *T. rangeli*. PCA revealed differential protein expression profiles based on MS1, iBAQ and LFQ features, illustrating the discrimination of *T. cruzi* DTUs from *T. cruzi marinkellei, T. dionisii, T. erneyi* and the generalist *T. rangeli*, included as an outgroup of the *Schizotrypanum* clade.

PCA of quantitative MS1 protein expression profiles clustered *T. c. marinkellei* closer to *T. cruzi cruzi* clade compared to the other bat trypanosomes such as *T. dionisii* and *T. erneyi*, and *T. rangeli*. Full genome analyses of generalist *T. c. cruzi* and *T. c. marinkellei* of bats have revealed that they share majority of the core genes, and that *T. c. marinkellei* is also able to invade non-bat cells (96), including human cells, *in vitro*. However, although *T. c. marinkellei* was never isolated from other hosts than bats, differences among the two genomes were mainly a smaller *T. c. marinkellei* genome compared to *T. cruzi* due to reduced number of repetitive sequences, with overall small nucleotide polymorphisms, and just one *T. c. marinkellei* specific gene, an acetyltransferase, which was identified in *T. c. marinkellei* and absent in *T. cruzi* Sylvio X10 and CL Brener genomes (96). Indeed, *T. c. marinkellei* has been included in many phylogenetic studies as an outgroup to *T. cruzi* DTUs (97, 98). Phylogenies inferred from multiple genes on bat trypanosomes clustered *T. c. cruzi, T. c. marinkellei* and the cosmopolitan *T. dionisii* in the *Schyzotrypanum* subgenus (99), which also included *T. erneyi*, a bat restricted trypanosome from African bats placed sister to *T. c. marinkellei* (52). *T. rangeli*, included as the outgroup, has a wide range of mammalian hosts, which are shared with *T. cruzi* including man, but are not pathogenic to mammalian hosts. Mixed infections with *T. cruzi* have been reported (100). *T. rangeli* belongs to the Tra[Tve-Tco] lineage, and is more closely related to Old World *T. vespertilionis* of bats and *T. conorhini* of rats than to the *Schizotrypanum* species (53). Based on our study, PCA analysis based on quantitative protein expression data sets from MS1, iBAQ and LFQ quantitative features showed congruence to well established evolutionary distances between *T. c. cruzi, T. c. marinkellei, T. dionisii, T. erneyi*, and *T. rangeli*.

Trypanosomes in the *Schizotrypanum* clade are confined to the digestive tract, where epimastigotes replicate in the midgut, and differentiate into metacyclic trypomastigotes in the hindgut from where they are passed through feces to mammalian hosts during a blood meal. On the contrary, *T. rangeli* is a Salivarian trypanosome which passes from the gut to the salivary haemocoel, and are passed to the mammalian hosts by bite during a blood meal. The identification of differentially regulated proteins between Stercorarian (*T. cruzi*) and Salivarian (*T. rangeli*) in the invertebrate hosts could help elucidate adaptations of the parasites to survive and proliferate in the hindgut or salivary glands. Studies have shown that different strains of *T. cruzi* result in varying outcomes of its infected invertebrate vectors. Peterson and colleagues (101) showed the significant variations in *Rhodnius prolixus’* survival and development following infections with different *T. cruzi* strains from DTU I. Differential protein expression between Stercorarian and Salivarian trypanosomes forms a platform to better understand the molecular interactions between the parasites and their invertebrate hosts.

Evolutionary relationships based on all three quantitative mass spectrometry features showed that *T. erneyi, T. dionisii* and *T. c. marinkellei* diverged earlier compared to *T. cruzi. T. c. marinkellei* formed the basal clade to *T. cruzi DTUs*, supported by high bootstrap support values in all analyses. High levels of correlation between phyloquant and phylogenetics were evidenced by Mantel test, and are in agreement with findings by Telleria *et al*., 2011 (49) who demonstrated the high correlation between gene expression (as shown by proteomic analysis using 2D-DIGE coupled to mass spectrometry) and phylogenetic diversity assayed by multilocus enzyme electrophoresis. Our study demonstrates the potential application of quantitative proteomics as an alternative method for taxonomic assignment of *T. cruzi* DTUs and closely related trypanosomes, and provides a valuable link between phylogenetic information and large-scale protein expression in the highly diverse *T. cruzi* strains.

The clustering of all replicates in the same monophyletic clades highlighted the reproducibility of our procedure. The evolutionary relationships of *T. cruzi* strains and closely related species based on LFQ values showed the highest correlation to evolutionary relationships of *T. cruzi* from phylogenies inferred by multilocus sequence typing, the golden standard proposed for the genotyping of *T. cruzi* in population studies (30). Using 32 different gene loci, both separately and in concatenation, Flores-Lopez and Machado (2011) reported the paraphyletic nature of *T. cruzi* II strains (now TcII-TcVI) (45), which were initially considered to be monophyletic. Based on MS1, iBAQ and LFQ intensities, our analyses are in agreement with the paraphyletic clustering of *T. cruzi* TcII-TcVI. Two major clades of *T. cruzi* were described by Flores-Lopez and Machado (2011); one with TcI, TcIV, TcIII, and one haplotype from TcV and TcVI; and the second clade haboring TcII, and the other haplotype from TcV and TcVI. In cases with haplotype loci for TcV and TcVI, three tree topologies were reported. In our analyses, iBAQ based clustering based on statistically significant values showed a topology with two major groups; one composed of TcIV, TcI-Tcbat sister clades, and TcIII, and the second clade composed of TcII, TcV and TcVI.

Two hypotheses have been proposed to explain the evolutionary relationships of *T. cruzi* strains and the hybrid lineages, TcV and TcVI (102, 103). Based on our analyses, the clustering of the two TcII strains, *T. cruzi Esmeraldo* and *T. cruzi* Y, with TcV and TcVI clades, respectively, demonstrated the parental and hybrid nature of TcII and TcV-TcVI, respectively. TcIII strains, though to be the other parental genotype, was basal to parental (TcII) and hybrid (TcV and TcVI) genotypes. The placement of DTU TcIV basal to the *T. cruzi* DTUs is supported by phylogenies based on different genetic markers, and a study by Flores-Lopez and colleagues (2011) using 22 concatenated nuclear loci demonstrated that TcIV was the earliest DTU to diverge from the common ancestor of *T. cruzi*, approximately 2.5 million years ago (45). In addition, phylogenetic analysis based on satellite DNA sequences corroborated that TcIV had an independent origin from the remaining DTUs (104). Based on protein expression level based on MS1 and statistically significant LFQ, TcIV and TcIII do not appear to be hybrid strains from TcI and TcII. However, based on iBAQ values, TcIV, Tcl-Tcbat and TcIII share a common recent ancestor. The close relationship of TcI, TcIII and TcIV has been proposed in the ‘Two hybridization’ model in which TcI and TcII hybridized to form the hybrid TcI-TcII, which mutated to form TcIII and TcIV. Contributing to clarifying the evolutionary history of DTUs, TcIII was not confirmed as the result of a hybridization event between TcI and TcII in analysis of satellite DNA (104).

Using large scale quantitative proteomics to map protein expression offers the opportunity to include in a phylogenetic clustering different protein features. This implies that genes exposed or not to selective pressure are jointly considered. Since differentially expressed proteins can be associated to genes exposed to selective pressure, and consequently poor selective markers, using total quantitative proteomic features should leverage this issue. However, using significantly regulated protein features produced well resolved evolutionary trees compared to the total quantified proteins (**Supplementary Figure 2**). Indeed, this method offers the possibility to identify strain/species specific quantitative markers for discrimination. In our analyses, a total of 3907 quantitative protein expression values based on LFQ features were used to classify the different strains/DTUs/species. Within this dataset, 2095 were regulated, while 1,812 were not differentially regulated (approximately 46%), meaning that the differentially expressed protein pool resolves the differences between the strains, while the not regulated proteins keeps the phylogenetic profile tight.

In this study a mass spectrometry data dependent acquisition method was used. In this method, precursor ions are selected in a stochastic manner creating missing values between replicates and samples. A way to overcome this limitation might be the use of data independent acquisition where all precursor ions within a certain m/z range will be selected and fragmented having a deeper coverage with less missing values. The method has the potential to be applied to different organisms improving our understanding on their evolutionary relationships.

## Conclusion

A novel approach to profile and infer evolutionary relationships in T. cruzi and close related trypanosome species based on large scale quantitative mass spectrometry analysis was described. This method enabled the clustering of T. cruzi strains to their DTUs, and showed close correlation to well established phylogenetic clustering.

Moreover, this approach allowed the identification of differentially regulated and DTU/ species-specific proteins. Differential protein expression profiles reveal upregulated or downregulated biological process and protein complexes between the DTUs and species, enhancing our knowledge on novel molecular basis of *T. cruzi* DTU diversity. In addition, DTU specific proteins offer potential application in the identification of DTU specific biomarkers, therapeutic targets and vaccine candidates.

Finally, we describe the expression of trypanosome multi-gene family proteins evolution during the trypanosomes’ divergence from their common ancestor. Using Pearson correlation, we show proteins whose expression increased, decreased or did not change with time, using *T. rangeli* as the common ancestor of the *Schizotrypanum* subgenus.

## Supplementary Figures

**Supplementary Figure 1.**
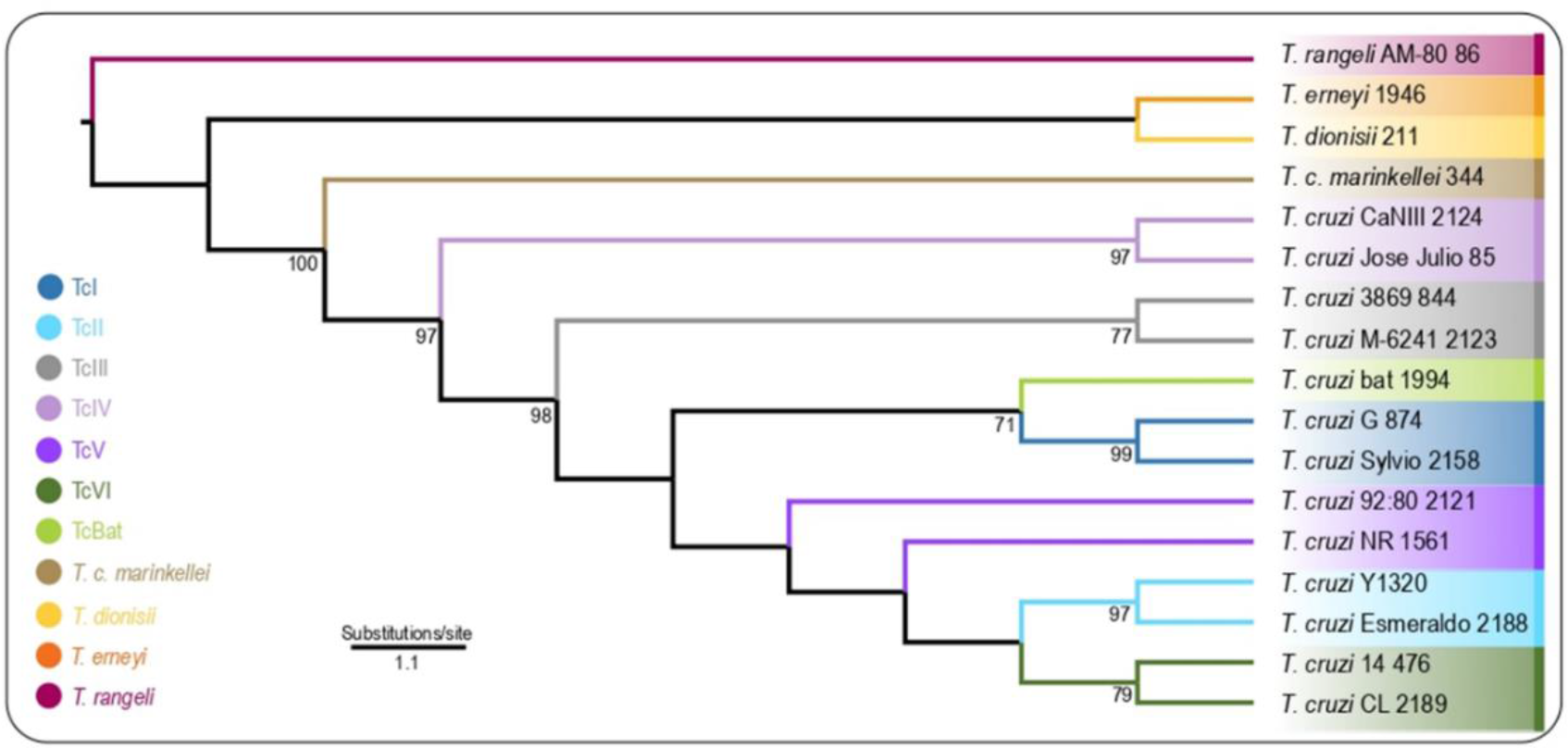
Phylogenetic tree showing clustering of *T. cruzi* DTUs and allied trypanosome species in the *Schizotrypanum* clade constructed using Maximum Parsimony phylogenetic algorithm. *T. rangeli* was included as an outgroup to the *Schzitrypanum* clade. Branch support values above 70 % are included.

**Supplementary figure 2:**
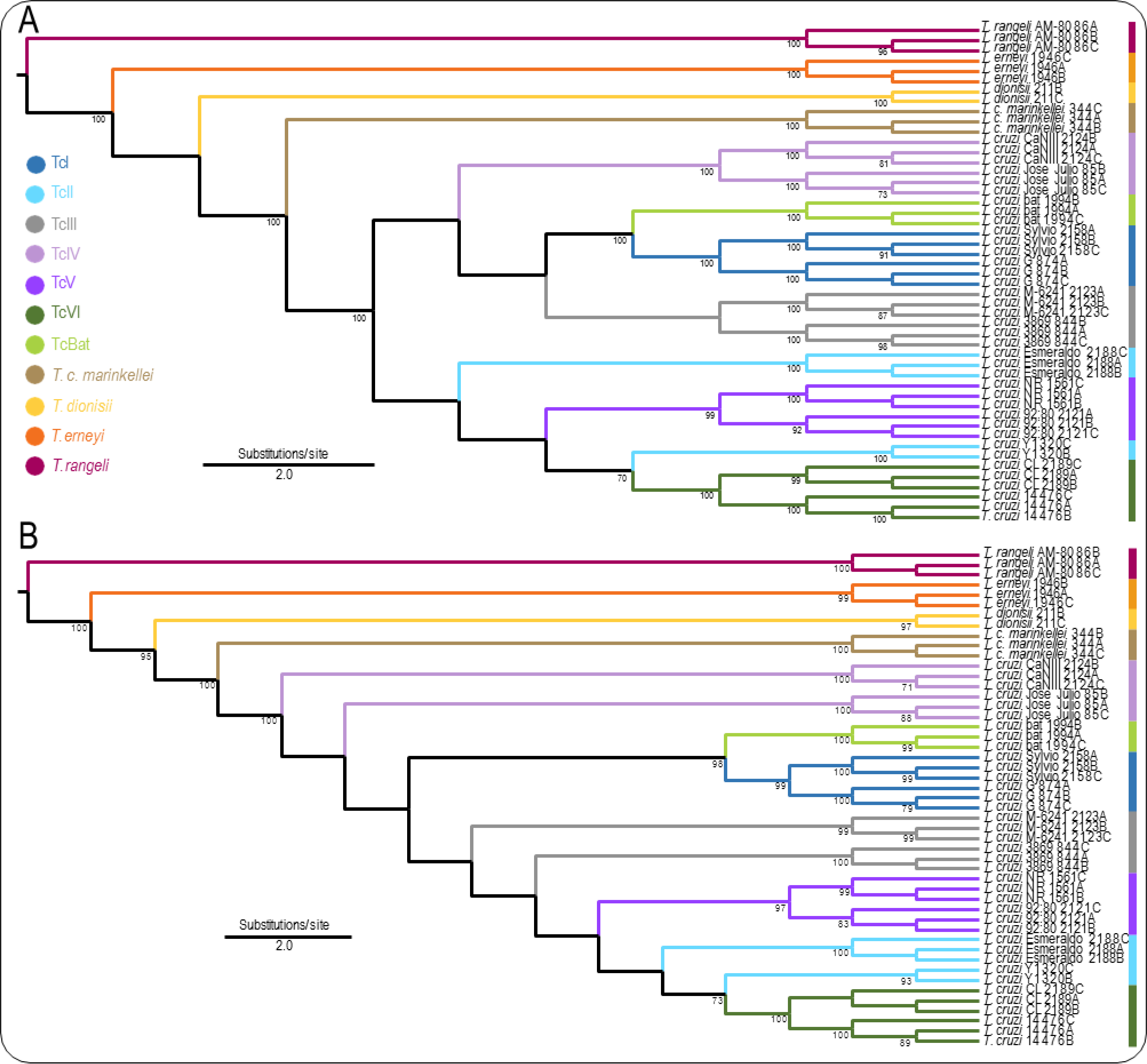
Phyloquant phylogeny based on total (A) iBAQ and (B) LFF values. Branch support values above 70 % are included.

**Supplementary Figure 3:**
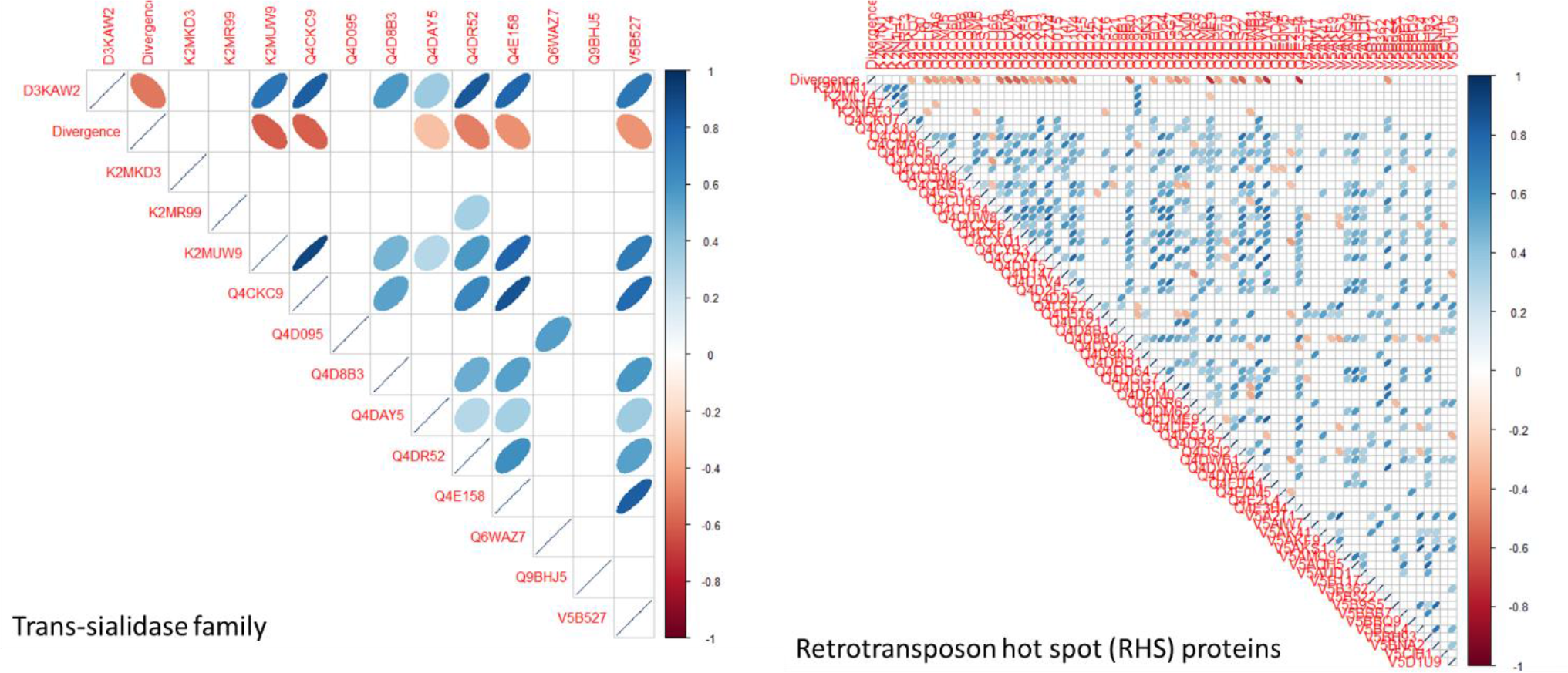
Correlograms showing Pearson correlation of selected multi-gene families (trans-sialidases and retrotransposon hot spot proteins) of trypanosome protein expression and divergence. Statistically significant correlations at a p-value < 0.05 are shown. Blue eclipses indicate positive correlation, while red eclipses indicate negative correlation from late to early appearing *T. cruzi* and closely related trypanosomes.

**Supplementary Figure 4:**
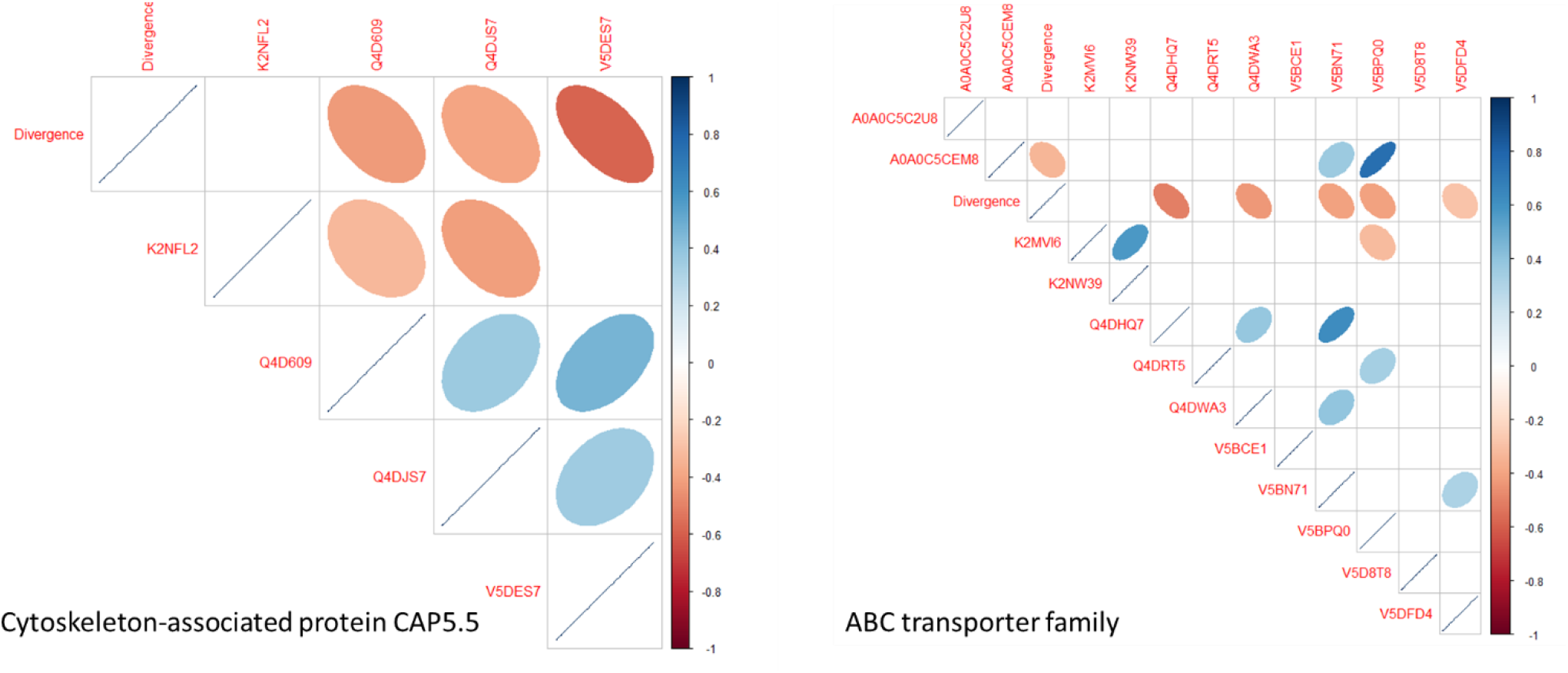
Correlograms showing Pearson correlation of selected multi-gene families (cytoskeleton-associated protein CAP5.5 and ABC transporter family) of trypanosome protein expression and divergence. Statistically significant correlations at a p-value < 0.05 are shown. Blue eclipses indicate positive correlation, while red eclipses indicate negative correlation from late to early appearing *T. cruzi* and closely related trypanosomes.

**Supplementary Figure 5:**
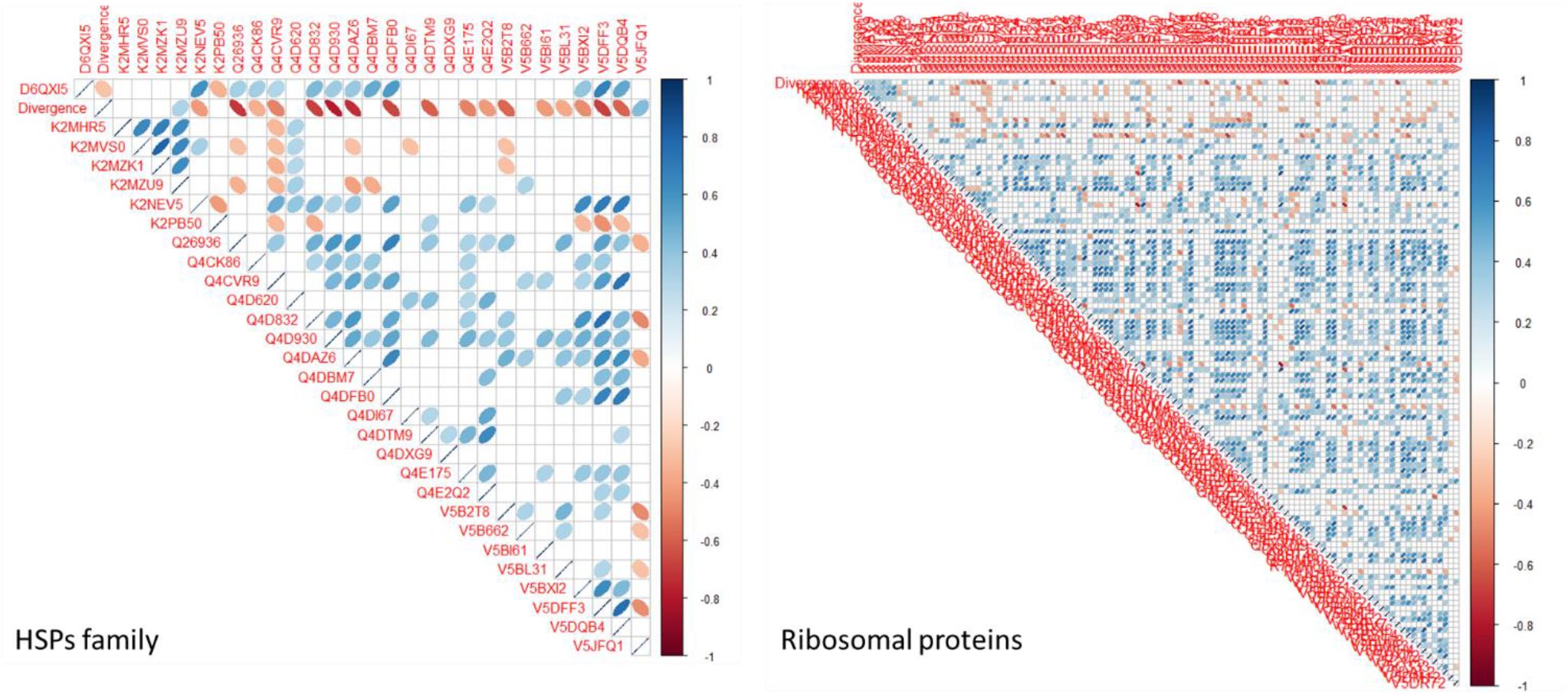
Correlograms showing Pearson correlation of selected multi-gene families (heat shock proteins family and ribosomal proteins) of trypanosome protein expression and divergence. Statistically significant correlations at a p-value < 0.05 are shown. Blue eclipses indicate positive correlation, while red eclipses indicate negative correlation from late to early appearing *T. cruzi* and closely related trypanosomes.

**Supplementary Figure 6:**
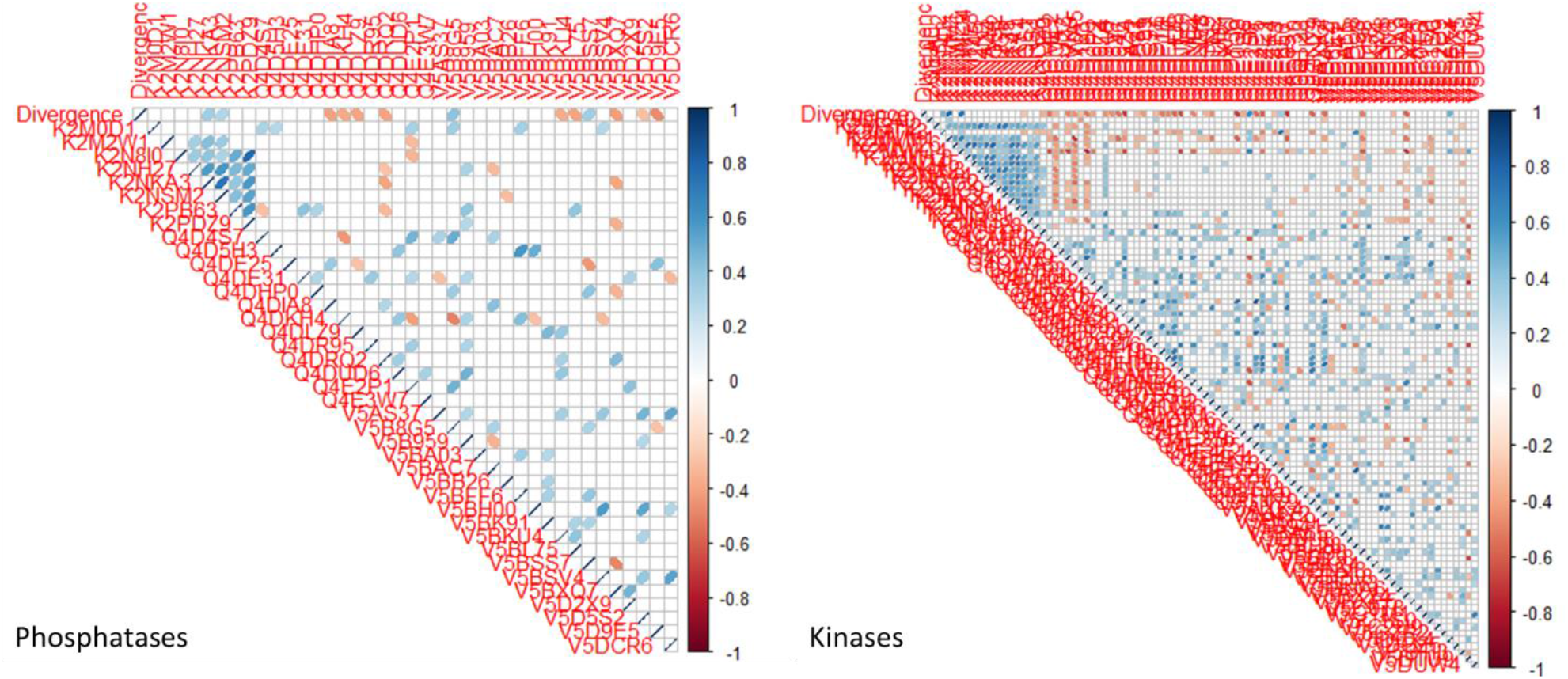
Correlograms showing Pearson correlation of selected multi-gene families (phosphatases and kinases) of trypanosome protein expression and divergence. Statistically significant correlations at a p-value < 0.05 are shown. Blue eclipses indicate positive correlation, while red eclipses indicate negative correlation from late to early appearing *T. cruzi* and closely related trypanosomes.

